# Fast prediction in marmoset reach-to-grasp movements for dynamic prey

**DOI:** 10.1101/2022.10.08.511417

**Authors:** Luke Shaw, Kuan Hong Wang, Jude Mitchell

## Abstract

Primates have evolved sophisticated visually guided reaching behaviors for interacting with dynamic objects, such as insects during foraging(P. S. Archambault, Ferrari-Toniolo, & Battaglia-Mayer, 2011; Bicca-Marques, 1999; Ngo et al., 2022; Smith & Smith, 2013; Sustaita et al., 2013). Reaching control in dynamic natural conditions requires active prediction of the target’s future position to compensate for visuo-motor processing delays and enhance online movement adjustments(Catania, 2009; Desmurget & Grafton, 2000; Fujioka, Aihara, Sumiya, Aihara, & Hiryu, 2016; Merchant & Georgopoulos, 2006; Mischiati et al., 2015; R. Shadmehr, Smith, & Krakauer, 2010; Wolpert & Kawato, 1998). Past reaching research in non-human primates mainly focused on seated subjects engaged in repeated ballistic arm movements to either stationary targets, or targets that instantaneously change position during the movement(Philippe S. Archambault, Caminiti, & Battaglia-Mayer, 2009; Battaglia-Mayer et al., 2013; Dickey, Amit, & Hatsopoulos, 2013; Georgopoulos, Kalaska, Caminiti, & Massey, 1983; Georgopoulos, Kalaska, & Massey, 1981). However, those approaches impose task constraints that limit the natural dynamics of reaching. A recent field study in marmoset monkeys highlights predictive aspects of visually-guided reaching during insect prey capture among wild marmoset monkeys(Ngo et al., 2022). To examine the complementary dynamics of similar natural behavior within a laboratory context we developed an ecologically motivated unrestrained reach-to-grasp task involving live crickets. We used multiple high-speed video cameras to capture the movements of marmosets and crickets stereoscopically and applied machine vision algorithms for marker-free object and hand tracking. Contrary to estimates under traditional constrained reaching paradigms, we find that reaching for dynamic targets can operate at incredibly short visuo-motor delays around 80 milliseconds, rivaling the speeds that are typical of the oculomotor systems during closed-loop visual pursuit(Cloherty, Yates, Graf, DeAngelis, & Mitchell, 2020). Multivariate linear regression modeling of the kinematic relationships between the hand and cricket velocity revealed that predictions of the expected future location can compensate for visuo-motor delays during fast reaching. These results suggest a critical role of visual prediction facilitating online movement adjustments for dynamic prey.

## Results

### Tracking of marmoset reaches

Primates exhibit a sophisticated repertoire of visually-guided reaching behaviors that enable them to interact with objects and navigate arboreal environments. While the motor aspects of these reaching movements are largely conserved across mammalian species, primates are unique in their integration of visual information to guide reaching movements(De Bruin, Sacrey, Brown, Doan, & Whishaw, 2008; J. H. Kaas, Gharbawie, & Stepniewska, 2011; Jon H. Kaas & Stepniewska, 2016; Sacrey & Whishaw, 2012; Wang et al., 2017). A recent study used video in the field to examine reach-to-grasp movements for flying insects among wild marmoset monkeys(Ngo et al., 2022). They found evidence to support an online active vision strategy during prey capture in which the head angle of marmosets actively tracked insects during reaching. However, the video resolution in the wild does not afford enough precision to measure visuo-motor delay nor test if the motor control strategy exploits predictions of future target location to compensate for processing delays. Here we establish an ecologically motivated reaching task within a laboratory setting using high-speed video to test the kinematics of reach-to-grasp movements for dynamic prey.

We used high-speed, multi-camera video and machine learning methods to track marmoset reaches for live, moving crickets. Reaching experiments were conducted by docking a mobile video recording platform to marmoset family cages within the marmoset colony and recording reaches using 3 GoPro cameras **(Figure 1A-B)**. Hands, fingers, crickets, and the experimental apparatus were tracked from three cameras using DeepLabCut(Mathis et al., 2018) and 3D position triangulated using Anipose(Karashchuk et al., 2021) (**Figure 1C-G)**. DeepLabCut performance using the labels illustrated in **Figure 1C-D** is quantified in **Supplemental Figure 1**. Although we were able to reconstruct 3D trajectories of reaching, most subsequent analyses focused on 2D tracking from the top view camera angle depicted in **Figure 1F**, because crickets mainly limited their motion to the horizontal plane during the reaching tasks.

**Figure 1.**
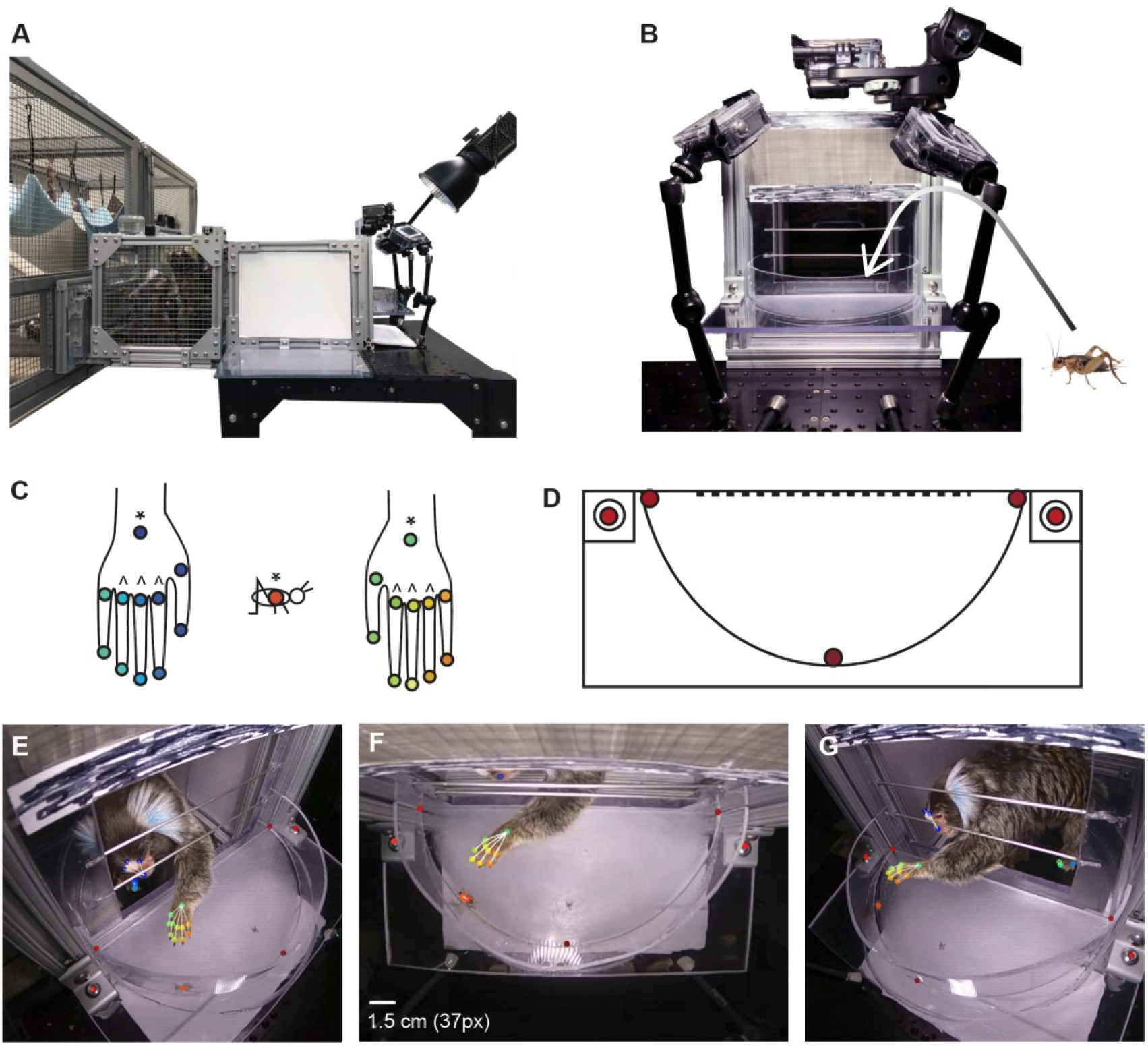
High-speed, multi-camera video reconstructs reaching for dynamic prey. **A:** A wheeled video platform allowed tests of individual marmosets from inside their housing in a colony. **B:** Three GoPro cameras were calibrated around a semi-circular reaching arena where a live cricket was placed. Marmosets reached through a rectangular aperture to acquire crickets. **C:** Feature points were manually labeled in a subset of video frames to train DeepLabCut. Hand and cricket markers (*) were used for reaching analyses and knuckle markers (^) for grasping analyses. **D:** Arena markers were used to validate spatial coordinates and correct for any post-calibration camera movement. **E-G:** DeepLabCut labeling of synchronized frames of video from three camera angles. DeepLabCut tracked labeled points with a RMSE of ∼3 and ∼7 pixels for the cricket and hand, respectively, compared to ∼37 pixels for the size of the marmoset wrist.

### Marmoset reach-to-grasp dynamics

Earlier work in laboratory settings have shown that marmosets are equally likely to prefer using their left or right hands in reaching(Hook & Rogers, 2008), and typically employ a power grasp strategy to reach items in which all fingers open in unison and close around the target to press it within the palm of the hand, which has been shown in marmoset (Fox, Mundinano, & Bourne, 2019) and macaque(Castiello & Dadda, 2019; Sartori, Camperio Ciani, Bulgheroni, & Castiello, 2013). A key difference from macaques is that marmosets do not use a precision grip. Old World primates have opposable thumbs that allow them to grasp small objects between the thumb and fingers (Marzke, 1997), while most New World primates, including the marmoset, use a power grasp(Bishop, 1962). We found that marmosets in our experiments predominantly employed a left or right single-handed (LH or RH) reaching strategy akin to a pause and lunge approach utilized by marmosets in the wild(Ngo et al., 2022; Schiel & Souto, 2017). This strategy may have been favored due to the reaching aperture utilized in our setup, which prevented marmosets from using their mouths or a two-handed pounce (P) strategy. We recorded a total of 271 reaches for live crickets (137 LH, 134 RH, 10 P) from 10 marmosets. Gross quantification of these reaches through a left-handedness index (LH reaches/total reaches) indicates an evenly split mixed hand preference over the population of reaches. Assessing this quantification at the marmoset participant level shows two marmosets with a strong right-handed preference (LHI=.31 and .04) and three marmosets with a strong left-handed preference (.83, .87, and .92), while the other 5 marmosets had little preference (.60, .68, .64, .66, .44, .40). Of the total recorded set of 271 reaches for crickets, 78 reaches contained crickets that moved during or prior to the reach, which were selected for subsequent analyses (see **Star Methods** for inclusion criteria).

Marmosets performed reach-to-grasp movements for moving crickets using a power grasp to enclose crickets within their hand at the end of the reach (**Figure 2**). In an individual reaching trial, the opening of grasp aperture begins as hand velocity drops with increasing proximity to target (**Figure 2A-C**), which is consistent with findings in previous NHP studies from both marmosets and macaques (Castiello & Dadda, 2019; Fox et al., 2019; Sartori et al., 2013). Finger separation distance was calculated as the average of the distances between the index finger and middle finger, and the middle finger and ring finger, using knuckle markers (**Figure 2B, dark blue points**). For this reach, there is a clear temporal order with the peak in hand speed followed by an increase in finger spread as the hand speed decreases (**Figure 2C**). This pattern was typical of the population of reaches. A frontal view of a typical reach (**Supplemental Figure 2**) shows the fingers wrapping around the cricket.

**Figure 2.**
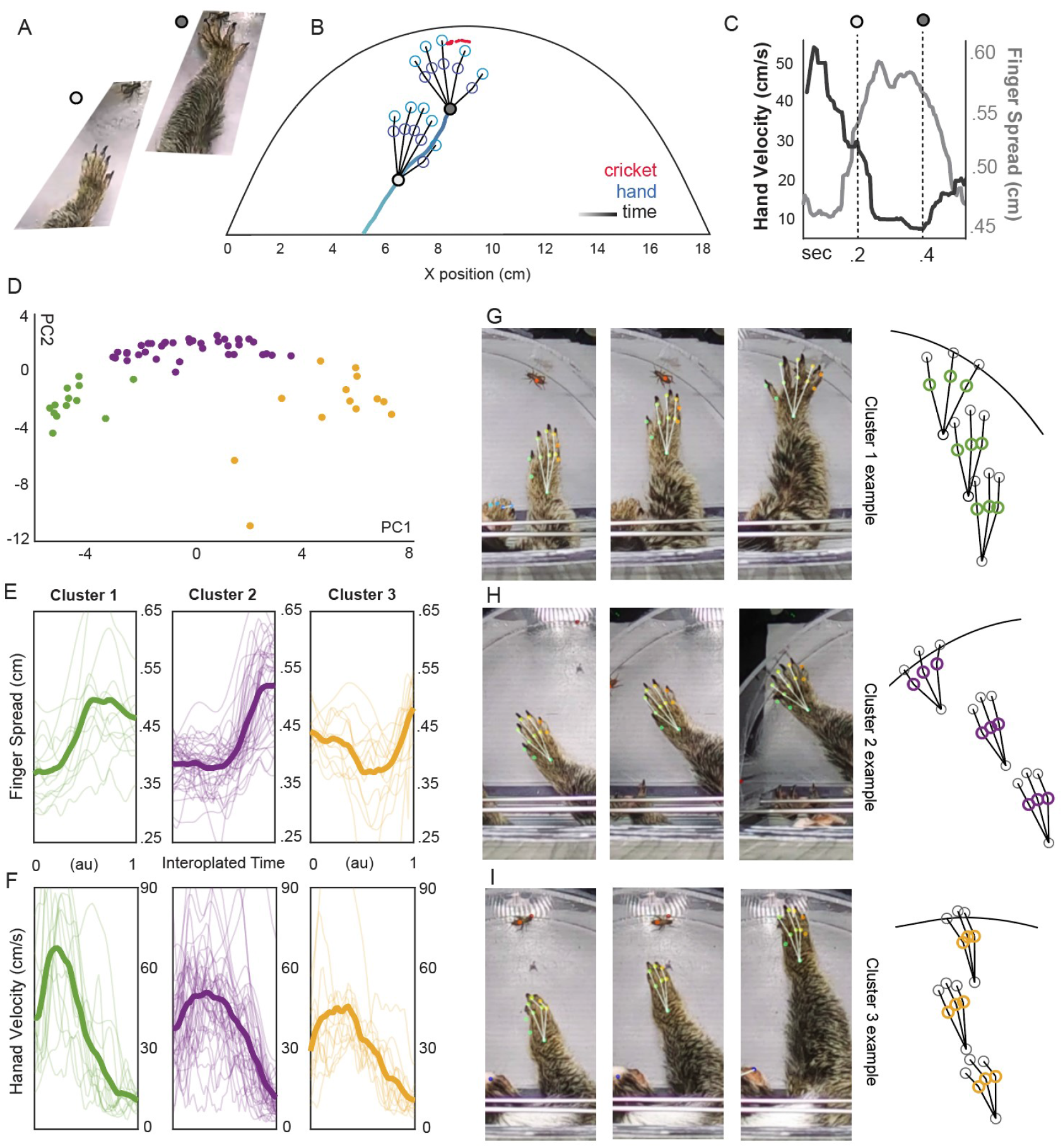
Hand aperture opens earlier with increasing hand speed in a power grasp. **A**: Cropped video frames show a single reaching trial at two time points indicated by light and dark gray points. **B:** The hand position for the corresponding time points in A is shown from the top camera view. The underlying light blue trace shows the trajectory of the reach from the central hand marker and the red traces indicates cricket motion. **C:** Within the example reach the speed of the hand (shown in black, left vertical scale) peaked and began to fall before the finger spread increased, depicted in gray (right vertical scale). **D:** The finger spread over time was submitted to a cluster analysis (see methods) which revealed a continuum along the principal component axes that we split into three clusters (green, purple, and orange) for visualization. The two outliers present in Cluster 3 (yellow, at the bottom) were due to finger contact against the retaining wall at the end of the reach. **E:** Finger separation peaked at longer delays in the reach for the three identified clusters when plotted over normalized time from reach start to cricket occlusion. **F:** The corresponding hand velocity traces for the same clusters reveal slower peak velocities correlating with later grasp opening. **G-I:** Three frames from an example reach to grasp for each of the clusters reveals a power grasp enclosing the cricket; on the right, plots of the three knuckle markers used to measure finger spread at corresponding times.

We used dimensionality reduction methods to further characterize the grasping strategies employed in the population of reaches. For each reach, a time series of finger spread distance was calculated based on the knuckle markers. The distance values were interpolated over a fixed number of timesteps over the total reach duration, with the reach start defined from the moment the hand first entered the arena to the end being when the fingers first began to close around the cricket. Principal component analysis of these reach trials captured 69.1% of the variance within the first two components. We found a continuum of points, where each point corresponds to one time-normalized reach, that extended along the first principal component as opposed to discrete clusters (**Figure 2D**). To visualize the variations in finger separation across the population of reaches we performed a k-means clustering and split the space into the best fitting three clusters.

The power grasp strategy shows a temporal pattern of earlier grasp opening for faster movements. Cluster 1 shows an early finger spread after reach initiation, which continues throughout the reach (**Figure 2E1**). By comparison in Cluster 2, the finger spread initiates later during hand transport (**Figure 2E2**), and in Cluster 3 finger spread initiates only at the very end of the reach **(Figure 2E3**). Plotting hand velocity time series for the same clusters we find higher and earlier peak velocities for Cluster 1 compared to Cluster 2, and likewise for Cluster 2 to 3 (**Figure 2F1-3**). Thus, the timing of grasp opening varies with reach speed. Video frames taken from an example reach in each of the three clusters illustrate the power grasp, with variation in the timing of grasp opening across the examples **(Figure 2G-I)**. In the third case with the slowest speed the fingers have still not fully separated by movement end (**Figure 2I)**. In contrast, rodents often use an arpeggio grasp to first contact the object with fingers extended flatly, and then based on somatosensory feedback, close around it(Whishaw & Gorny, 1994). Here even the slowest reaches appear to involve opening of the grasp near contact (**Fig. 2E3**), and thus are more consistent with a vision-guided power grasp strategy typical of macaque reaching.

### Hand velocity converges to match lateral cricket velocity during reaching

Movement dynamics during prey capture have been found to follow control strategies for target interception such as pure pursuit and proportional navigation across a range of species including dragonflies(Mischiati et al., 2015), hawks(Brighton & Taylor, 2019), and humans(Lenoir, Musch, Janssens, Thiery, & Uyttenhove, 1999). In pure pursuit the rotational acceleration, or steering, of the body (or hand) is dictated by the instantaneous offset from the target location, called the range vector. Pure pursuit seeks to close the distance of the range vector but ignores motion of the target and is not anticipatory of its future location. By contrast, proportional navigation aims to maintain a constant bearing angle of the range vector as the pursuer closes the distance. One way to visualize the distinction between these strategies is to decompose velocity into a component that lies along the range vector at each movement, the direct axis, and another component that lies on its orthogonal projection, the lateral axis. Both strategies predict changes in velocity related to closing distance on the direct axis, but in addition proportional navigation predicts a matching of the velocity of the body (or hand) with the lateral velocity of the target to maintain a constant bearing angle. Another feature characteristic of proportional navigation is that the derivative of the range vector over time will be negatively correlated at −1 with the range vector itself(Mischiati et al., 2015).

We examined the direct and lateral components of hand velocity as marmosets performed reaches to moving crickets in the arena (**Figure 3)**. Although we are able to reconstruct the full 3D kinematics of reaching movements **(Figure 3A)**, we focus our subsequent analyses on the horizontal plane seen from the top camera view which captured most of the cricket motion. To examine the influence of cricket velocity on hand velocity, we decomposed velocity of the hand (H) and cricket (C) into direct and lateral components relative to the range vector (RV) connecting the instantaneous position of the two **(Figure 3B)**. In the set of reaches, most of the hand motion falls along the direct axis, *Dh*, which is necessary to traverse the arena to close on the cricket and exhibits a peaked shape typical of ballistic reaching movements (**Figure 3C**, dashed lines). Conversely, most of cricket motion falls along the lateral axis, *Lc*, due to the tendency of crickets to crawl along the perimeter of the reaching arena. The lateral component of absolute hand velocity, *Lh*, was closely matched in magnitude to cricket lateral velocity, *Lc* (**Figure 3C**, solid lines). We thus sought to determine to what extent the lateral hand velocity might match cricket velocity, both in sign and magnitude over the reach, as predicted during proportional navigation.

**Figure 3.**
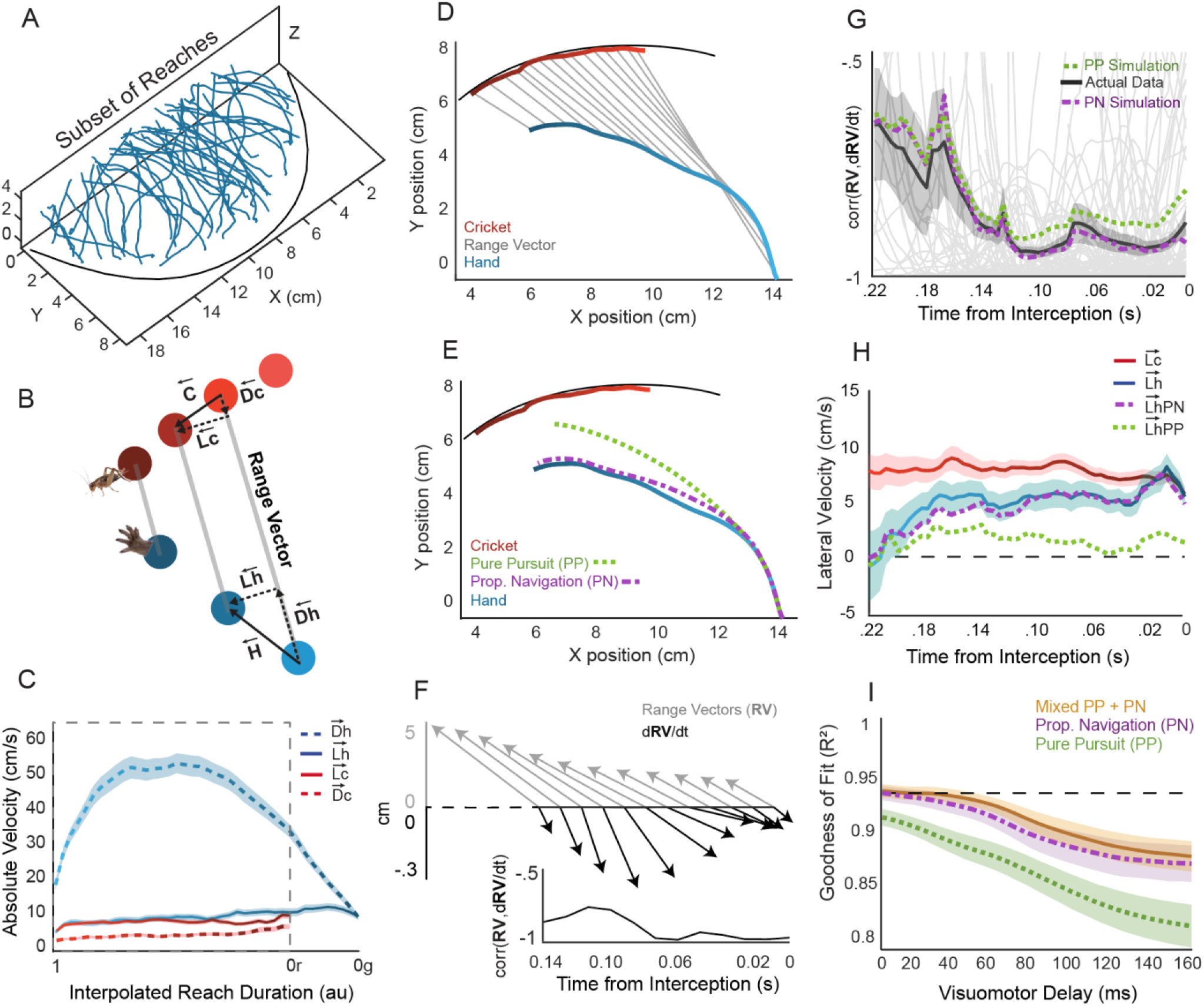
Hand velocity during reaching is consistent with proportional navigation. **A:** Representative 3D reconstructions f the population of reaches for moving crickets. **B:** At any moment, the hand and cricket velocity can be broken down into two omponents, the direct and lateral velocity. Direct velocity is the component projected along the range vector while lateral is he orthogonal projection. **C:** The average absolute cricket and hand velocity for the direct and lateral components as illustrated n **B** over interpolated time for the population of reaches. The dashed box indicates the period where the cricket was visible. **D:** ath of the hand (blue) and cricket (red) in an individual example reaching trial. Lighter points indicate the beginning of the each. Gray lines indicate line of sight, or range vectors, for corresponding hand and cricket points, 1:4 down-sampled. **E:** A ure pursuit (PP) steering strategy aligns hand velocity with the range vector. Proportional navigation (PN) rotates hand velocity o mirror range vector rotations. Overlaid simulations of PP and PN show that PN best approximates this reach. **F:** PN strategies redict matching the lateral component of the hand and cricket velocity while the range to the target is close, and if followed, he derivative of the range vector during pursuit should be in opposite direction of the range vector. For the example reach, the ange vectors are shown in gray (matching gray lines in **D**) and their derivatives in black, with the correlation between vectors ver time quantified below (black). The trace is aligned on the right to the moment of interception. **G:** The correlations between he range vectors and their derivatives converge towards −1 on average across the population of 78 reaches (individual reaches hown as gray lines, average in black, +/-1 SEM in gray shade). This analysis was also performed for PP and PN simulations, howing that PN is a better match to actual data. **H**: Lateral velocities from the hand and pursuit simulations were made positive hen traveling in the same direction as cricket velocity. Hand lateral velocity increasingly approaches cricket lateral velocity hroughout the reach. This is better matched by a PN simulation than a PP simulation. **I:** Pursuit strategy simulations perform orse when a delay is introduced. We include a mixed model simulation, which uses a combination of PN and PP to steer hand elocity and is moderately more robust than proportional navigation alone. Goodness of fit quantified by R^2^ values that compare model simulations to actual hand data. All error fields are +/-1 SEM.

A typical reaching trial demonstrates that hand velocity tracks the lateral motion of the cricket target in a manner more consistent with a proportional navigation strategy for interception **(Figure 3D-F)**. In this trial, the cricket moves in the horizontal plane along the boundary of the arena and the hand adopts a curved trajectory in pursuit (**Figure 3D**). We applied computational methods from a previous study of hawk prey pursuit(Brighton & Taylor, 2019) to simulate the hand’s trajectory using either a pure pursuit (PP) or proportional navigation (PN) steering model. For this trial the PN model provides a closer match to the actual hand trajectory (R^2^=.99) compared to the PP model (R^2^=.94) (**Figure 3E**). In line with the better fit of the PN model, the correlation of the range vectors to their derivatives over time for this example trial takes on values near −1, especially near the end of the reach as the hand converges upon the target. In pure pursuit, the lateral velocity is not necessarily matched, leading to overall weaker negative correlations. We find that the correlation increasingly converges towards −1 up to interception in this reach (**Figure 3F**).

When steering strategies are examined over the set of reaches for moving crickets, the PN model better explains the hand kinematics than the PP model. We applied PN and PP model simulations to the population of reaches and compared the range vector correlations of the models against the actual movements. We find that the mean correlation coefficient converges towards −1 over the duration of reaches, as in the previous example trial, and that the correlation trace of the PN steering model better matches that of the hand than the PP model, especially as the hand nears interception (**Figure 3G**). To examine if the lateral velocity of the cricket was matched by hand velocity, we examined the mean of the lateral velocity of the hand multiplied by the sign of instantaneous cricket velocity, such that correspondence would give similar mean velocity and no correlation in direction would converge towards zero. This process was repeated for the lateral velocity components of PP and PN simulations. The mean lateral velocity of the hand, *Lh*, increasingly matches that of the cricket, *Lc*, over the time-course of reaches, which is very closely approximated by the PN model, but not the PP model (**Figure 3H**). Together these results provide evidence to support a proportional navigation strategy of hand guidance. However, a weakness of any of these steering models stems from their ability to compensate for visual processing delays, which represent a critical feature of any real biological or engineered system. Indeed, when we incorporate a visual delay of cricket position into the PP and PN models we find the goodness of fit for simulated kinematics break down progressively with visuomotor delay **(Figure 3I)**. A previous study found that a mixed pursuit strategy, including an additive combination of PP and PN guidance commands, was more robust than single strategy models(Brighton & Taylor, 2019). However, even implementing a mixed PP+PN strategy here, we find that performance matching the hand kinematics breaks down progressively with delay. Therefore, we sought to determine visuo-motor delay in marmoset reaching for this task, and furthermore to test if an explicitly predictive control strategy would be able to better explain observed reach kinematics while using realistic delay values.

### Hand velocity incorporates target predictions to compensate for visuo-motor delays

A key problem in guiding the hand during pursuit is that positional and velocity information about the target arrives at a processing delay in the visual system before it can influence actions. Motor control theories often assume predictive models that estimate future body and target positions to compensate for such delays in planning movements(Reza Shadmehr & Krakauer, 2008; Wolpert & Kawato, 1998). In human experiments employing double-step reaching tasks, where an initial target turns off and a new illuminated target turns on mid-reach, there are visuo-motor delays ranging from 125-225 ms(Brenner & Smeets, 1997; Day & Lyon, 2000; De Brouwer & Spering, 2021; Prablanc & Martin, 1992). By contrast, studies with head-restrained macaques performing double-step tasks have found substantially longer motor delays around 200-300ms(Philippe S. Archambault et al., 2009; P. S. Archambault et al., 2011; Battaglia-Mayer et al., 2013; Dickey et al., 2013; Georgopoulos et al., 1981). This slower timing may reflect constraints due to head-restraint and the artificial nature of the tasks. Thus, it was critical to first asses the visuo-motor delays in the current task.

Visuo-motor delays in dynamic reaching for cricket prey were much shorter than those found in previous non-human primate studies, ranging between 80-100 ms. To estimate visuo-motor delay, we identified discrete cricket acceleration events across task trials in which the cricket increased its motion while the reach was ongoing. The lateral speed of the cricket was filtered to flag the top 5% of moments where lateral velocity increased. We observed that immediately following an increase in lateral cricket velocity, there is a corresponding increase in lateral hand velocity in the same direction that begins around 80 ms (**Figure 4A**). This analysis, however, relies on discrete events in the total data set. To determine if it was consistent throughout the entire data set, we applied a simple linear regression on all the velocity data to predict lateral hand-velocity based on the recent history of cricket lateral velocity (see **Star Methods, Equation 1**). This enabled us to fit a linear kernel for hand velocity based on cricket velocity (i.e., the impulse response function corrected for autocorrelations in cricket velocity). Consistent with the visuo-motor delay observed due to sudden jumps, the linear prediction found a significant prediction coefficient near 80ms (p<.0001 at 87.5ms and p=.0559 at 100ms), but not at other delays (**Figure 4B**). This suggests that changes in hand velocity track the cricket velocity at relatively short delays around 80-100ms, a value much faster than the 200-300ms reported in constrained tasks (Philippe S. Archambault et al., 2009; Battaglia-Mayer et al., 2013; Dickey et al., 2013; Georgopoulos et al., 1981). However, this value is within range of the delay seen in ocular following responses in marmosets, which is close to 75ms(Cloherty et al., 2020). Of interest, like the reaches for live crickets, studies of oculomotor pursuit employ fixation targets that are moving continuously whereas the double-step reaching studies in macaques used instantaneous jumps in target positions.

**Figure 4.**
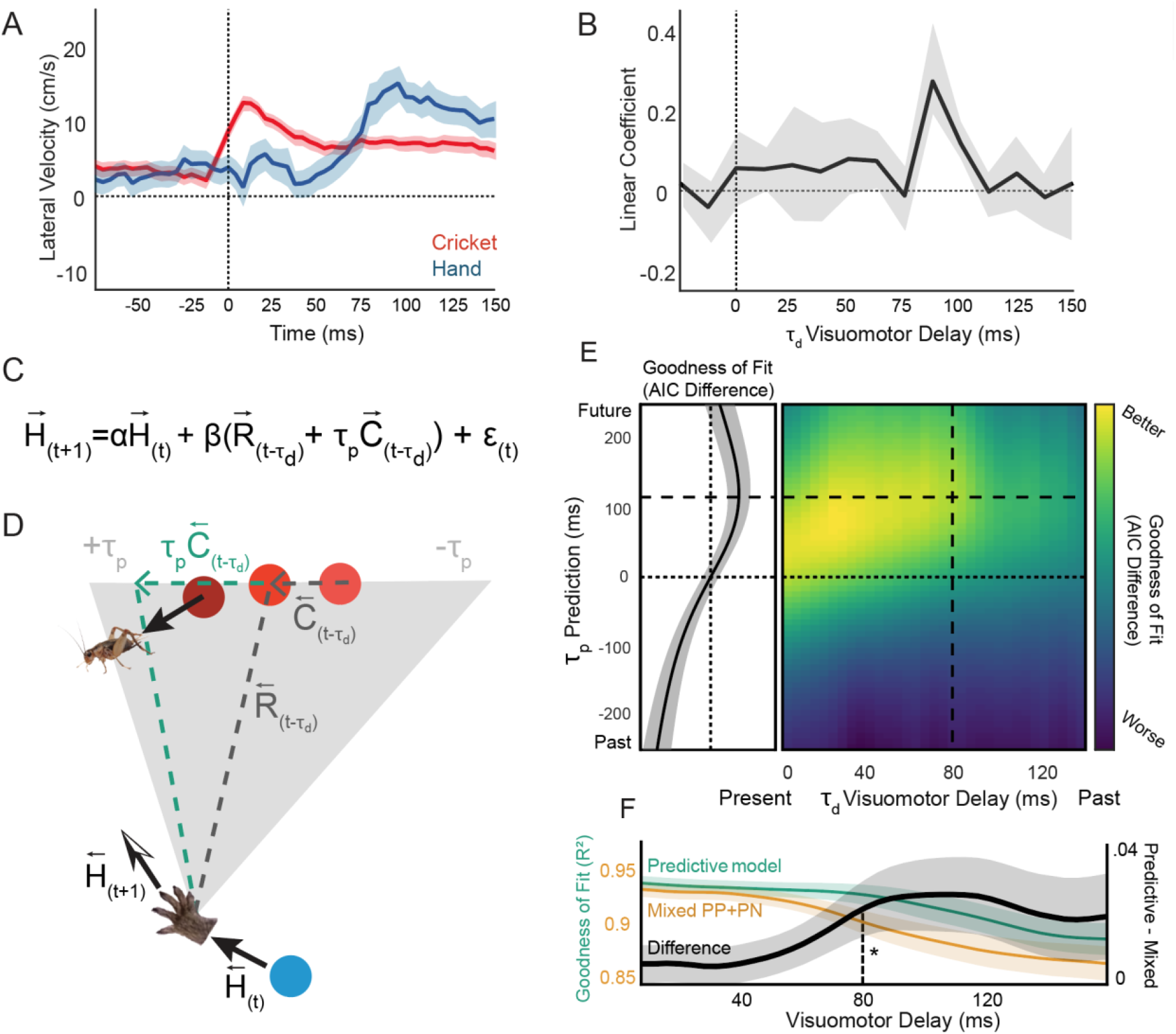
Cricket velocity predictively guides hand velocity to compensate for visuo-motor delay. **A:** Increases in the lateral cricket velocity are followed by increases in lateral hand velocity at a latency of roughly 80ms. A Difference of Gaussian filter was applied to the absolute velocity of cricket motion to identify discrete acceleration events (N=68). The average cricket velocity (in red) time-locked to those events reflects increases in speed at time zero, as expected, while hand velocity follows with a comparable acceleration at a delay (blue), shown as +/-1 SEM. **B**. A linear regression of lateral hand velocity with the offset lateral cricket velocity from delays ranging from −25 to 150 ms reveals a significant regression coefficient between 80-100 ms. Error shown in 1 SEM estimated from a jackknife procedure. **C:** The multivariate linear model equation used to test the influence of visuomotor delay τ _d_ and prediction time constant τ _p_ on the fit of hand velocity. **D:** The model estimates the hand velocity at the next time step (t+1) based on the range vector (R) at time t-τ_d_ and momentum from the previous hand velocity. A predictive term based on cricket velocity (C) at time t-τ _d_ (teal dashed lines) is multiplied by the prediction term τ _p_ to project the range vector into its estimated future location (teal dashed lines). The gray field represents the range of tested prediction τ _p_ values. **E:** The goodness of fit as a function of τ _d_ and τ _p_ is shown in color scale. Upon setting τ _d_ to 80ms, the best fit of this model is achieved using a τ_p_ prediction = 115ms (indicated by intersection of black dashed lines). Color scale indicates the goodness-of-fit surface measured by a difference of AIC values between the full predictive model (equation as in **C)** and a model that only uses hand velocity (momentum term) as a predictor. (Left panel) The goodness-of-fit with visuo-motor delay τ _d_ fixed at 80ms and varying the prediction time constant τ_p_ is shown with 1 SEM estimated from a jackknife procedure. **F:** When this predictive pursuit model is applied in an approach to simulate hand trajectories (as in Figure 3) it provides a better fit (teal curve) that degrades more gradually than the best mixed PP+PN steering models (orange curve, left vertical axis). Colored traces indicate mean R^2^ of simulation fits with 1 SEM fields. The model improvement for the predictive vs. best steering model (black curve, right vertical axis) at 80ms delay and beyond is significantly better for the predictive models (Sign Rank test p=.0105).

While the visuo-motor delays are briefer in the current task than previous studies, there may still be significant advantages for taking them into account by planning predictive movements that incorporate the cricket’s velocity. To test the role of prediction in guiding the hand, we fit an auto-regressive model of hand velocity that included terms for the visuo-motor delay, τ_d_, as well as a prediction time constant, τ _p_, that dictated how far into the future the cricket’s position was projected based on its velocity. We followed a generative model formulation from a previous study in macaques(Yoo, Tu, Piantadosi, & Hayden, 2020), which described the hand velocity in terms ofa momentum term plus a scaled version of a range vector, R(t-τ_d_), at some visuo-motor delay (**see equation, Figure 4C**). To compensate for delays in the range vector (**cyan dashed line, Figure 4D**) the model includes a term to estimate the cricket’s future position, τ_p_ C(t-τ_d_), where τ_p_ is a linear temporal weight that determines how far in the future to adjust for the cricket’s future position (**dark gray dashed line, Figure 4D**). We pooled data from 78 reaches for moving crickets and tested the predictive model fit for visuo-motor delays τ_d_ over a range from 0ms to 145ms in the past and prediction time constant τ_p_ over a range from 250ms in the past up to 250ms in the future. We assessed model fit by the difference of Akaike Information Criterion (AIC)(Akaike, 1973) values for the full model (including cricket velocity prediction) against a baseline model that only contains information about hand velocity (momentum term).

Auto-regressive modeling indicates that hand velocity is best explained by a model where visuo-motor delay is compensated for by using predictions based on cricket velocity. The AIC difference, or goodness-of-fit, between the full predictive model and the hand velocity model based on momentum showed the best fit with a positively correlated diagonal structure reflecting that positive temporal prediction constants were matched to compensate for visuomotor delays (**yellow band in Figure 4E**). When we set delay to 80ms as determined above (**Figure 4A, B**), the best fit is achieved using a prediction constant of 115 ms into the future, (**vertical dashed line, Fig. 4E**). To visualize standard error in these surface estimates, the cross-section at that 80ms visuo-motor delay was examined as a function of the temporal prediction parameter, with its confidence intervals estimated using a jackknife procedure (**left panel, Fig. 4E**). The prediction model demonstrated a significant improvement in model performance for temporal constants weighted into the future. To obtain a more direct comparison of this model against the PP, PN, and PP+PN mixed steering models **(Figure 3I)**, we tested this prediction model using a simulation approach directly comparable to earlier analyses fitting hand kinematics, and tested a range of visuo-motor delays. When compared against the best steering model, the PP+PN mixed strategy, the predictive model outperformed it at each delay, showing a more gradual breakdown in performance for longer visuo-motor delays **(Figure 4F, cyan versus orange curves)**. Taking the difference in performance between models, the full predictive pursuit model significantly outperformed the mixed PP+PN steering model at 80 ms (Sign Rank test, p = 0.0105) and longer delays **(Figure 4F, black trace)**. These findings support that a predictive term compensating for visuo-motor delay provides a better description of reaching kinematics at realistic delays. Furthermore, while steering models such as proportional navigation successfully describe marmoset reaching for moving crickets when they are formulated with no processing delays **(Figure 3)**, they lack the ability to do so given systematic delays characteristic of the real system.

## Discussion

The present findings demonstrate that during an ecologically motivated reaching paradigm for moving crickets, marmoset monkeys utilize a power grasp that opens earlier in the reach for faster velocity reaches. Furthermore, marmosets guide their hand movements at very brief visuo-motor delays, comparable to oculomotor pursuit. In an examination of hand guidance steering models, we find that proportional navigation explains hand motion reasonably well and provides better fits than a pure pursuit model (Figure 3). However, these models perform worse in the presence of realistic visuo-motor delays. To explain the reach kinematics given a realistic visuo-motor delay, which we estimate to be around 80 ms, we find that it is necessary to use a predictive pursuit model that incorporates a term to extrapolate the future location of the target based on cricket velocity. This predictive pursuit model outperforms the proportional navigation or mixed strategy model(Brighton & Taylor, 2019) at the estimated delay. We find that the prediction constant closely matches the scale of the visuo-motor delay, extending slightly beyond it about 35 ms into the future. Such a prediction constant, that remains close to or slightly beyond the visuo-motor delay, is often close to optimal for maintaining stability and achieving optimal solutions in Bayesian integration(Körding & Wolpert, 2004) or Smith prediction(Read et al., 2022).

The current finding emphasizes that moving towards realistic dynamics in experimental designs c(Parker, Abe, Leonard, Martins, & Niell, 2022)an better inform our understanding of reaching behavior, and provide insights into the underlying neural correlates and the evolutionary contexts that have driven their development. A previous study also found that primates use prediction in a pursuit task involving a simulated evading virtual target that was captured by a screen cursor under joystick control(Yoo et al., 2020). In our study we have adapted their approach to modeling prediction, but have done so within a task where reaching movements unfold at natural timescales. The former study identified predictive terms beyond 500 ms into the future, which match the delays imposed by using a joystick for control in their task. Here we find that similar predictive strategies are involved to compensate delays when reaching for live prey, but that they occur at much shorter latencies where visuo-motor delays are comparable to oculomotor pursuit and under 100 ms. While the present findings were sufficient to characterize the visuomotor strategies in the current reaching task, future studies could leverage recent advances in head mounted eye tracking(Meyer, Poort, O’Keefe, Sahani, & Linden, 2018; Parker et al., 2022) to characterize coordination between head and eye gaze in guiding predictive reaching.

Visually guided insect predation and fruit/leaf eating in an arboreal environment are hypothesized to play a role in the evolution of primates(Cartmill, 1974; Sussman, Tab Rasmussen, & Raven, 2013). Although not ubiquitous, insect hunting is not only observed in New World monkeys, like marmosets, but also in Old World monkeys and hominoids(McGrew, 2014; Raubenheimer & Rothman, 2013). Fronto-parietal neural circuits for motor control in primates are thought to have evolved in primates to optimize visuo-motor strategies for coordination and speed of reach-to-grasp movements(Karl & Whishaw, 2013). Much evidence suggests that motor commands are sent via feedback pathways from premotor cortex to higher order sensory areas in parietal cortex, and could mediate the forward models necessary to implement predictive on-line control(Reza Shadmehr & Krakauer, 2008; Wagner & Smith, 2008). The current experimental paradigm shows that predictive strategies in reaching emerge naturally even within a laboratory context for dynamic targets, such as crickets, affording new opportunities to study the underlying neural mechanisms within an ethologically motivated paradigm.

## Acknowledgments

We would like to thank Dina Graf and members of the Mitchell lab for help with marmoset care and handling. We also thank Cory Miller, Alex Huk, and Cris Niell for comments on this manuscript. This work was supported by NIH grant UF1NS116377 (JFM and LS) and a grant from the Schmitt Program on Integrative Neuroscience (KHW and JFM) and the Del Monte Institute for Neuroscience at the University of Rochester (KHW and LS).

## Author Contributions

Conceptualization, all authors; formal analysis, all authors; investigation, all authors; methodology, all authors; data acquisition, LS; writing – review & editing, all authors.

## Inclusion and Diversity

None of the authors of this paper self-identifies as an underrepresented ethnic minority in their field of research or within their geographical location. We support inclusive, diverse, and equitable conduct of research.

## STAR METHODS

### Resource Availability Lead Contact

Further information and requests for resources and reagents should be directed to and will be fulfilled by the lead contacts Jude Mitchell and Kuan Hong Wang.

### Materials Availability

This study did not generate new unique reagents.

### Data and Code Availability

All data reported in this paper will be shared by the lead contact upon request. All original code in this paper is available in Supplemental Materials.

## EXPERIMENTAL MODEL AND SUBJECT DETAILS

All experimental procedures described here are approved by the University Committee on Animal Resources at University of Rochester. 10 marmoset monkeys (5 females, 5 males) were recorded by multiple cameras while reaching for live crickets and cheerios. Their ages ranged from 1.2 years old to 7.1 years old. The crickets used in experiments served as additional enrichment to their daily diet.

## METHOD DETAILS

### Reaching Task and Video Recording

Reaching experiments were conducted by docking a mobile video recording platform to marmoset family cages within the marmoset colony and recording reaches using 3 GoPro Hero 7 Black cameras **(Figure 1B)**. Marmosets were acclimated to the experimental apparatus and allowed to become comfortable reaching for cheerios as food treats, and then also for crickets. Acclimation was typically less than 30 minutes and was necessary only on the first day of experiments. To calibrate cameras prior to reaching experiments, we recorded a brief video of an 8×6 checkerboard array that was moved and rotated continuously within the reach arena **(Figure 1B,D)**. Frames of this video were extracted and fed into triangulation algorithms provided in the Anipose toolkit(Karashchuk et al., 2021). A manually triggered photography flash aimed at the cameras was used to deliver a synchronization signal to all three cameras. This created 1-2 frames of video that were significantly over-exposed and easily identified using custom Matlab (Mathworks) and ffmpeg (Ffmpeg Developers) scripts.

Up to 15 cricket reaches were recorded from an individual marmoset on a given recording day. A sliding plate at the entry of the reaching enclosure was closed to isolate individual marmosets briefly from their family groups on the test platform. Reaching trials began by sliding an opaque acrylic plate upwards that occluded the view of the reaching platform, thus allowing the marmoset to view the target and begin hunting. The marmoset was able to reach through a large rectangular aperture (9cm x 13.5cm) that contained two laterally traversing 2mm stainless-steel bars at 3cm and 6cm height and was built into an acrylic plate. This aperture allowed for a wide range of reach start points while also preventing the animal from getting outside the test chamber **(Figure 1B)**. Before lifting the clear acrylic sheet to allow reaching to begin, the crickets were dropped onto the platform and allowed to move freely. Crickets were kept from crawling off the platform with a 4cm tall acrylic wall molded into an arc and cemented to the platform with a center maximum y-dimension of 9cm and base maximum x-dimension of 20cm. The platform was sanded to eliminate reflections and increase hand and target contrast. Reflection reduction was useful due to the competition of the features of a tracked subject’s reflection against the features of the subject itself.

Our video recordings were optimized to capture high velocity ballistic reaches. The GoPro Hero 7 Black was specifically chosen for its high-speed capture capability and manual control of recording settings. High frame rate (1/240s) and high shutter speeds (1/960s) were sufficient to render all hand features in full detail without observable motion blur. To attain proper exposure at this high shutter speed and to avoid shining a bright light directly into the eyes of our subjects, a LED light source (Nanlite Forza 60), was positioned to project light at a 65-degree angle 45cm from the reaching platform **(Figure 1B)**. Light intensity at the platform was measured to be 4.6E5 Lux, comparable to direct sun on a bright day. The cameras were arranged around the reaching arena to provide a view from above and a downward angled view from the right and left **(Figure 1E-G)**. The cameras were fixed in position using articulating camera brackets mounted to an aluminum breadboard.

### Video Preparation and DeepLabCut Pose Estimation

Videos were preprocessed by the experimenter to indicate which video frames contained reaches for crickets and to provide metadata for each reach. Reach success, handedness, and whether reaches were composed of a single volley or multiple volleys were documented for each reach in the data workflow. We defined volley multiplicity by whether the hand returned fully to the reach aperture opening on the platform before extending again. Selected frames spanned from ∼200ms prior to the hand moving through the reaching aperture to full retraction of the arm, which was typically accompanied by delivery of the target to the mouth. These data were used to generate standalone video files for each reach that were synchronized for all camera views. Adobe Premiere Pro was used for video viewing and scoring. Videos were cropped to 760×720 pixels to remove pixel areas with irrelevant information. DeepLabCut 2.1.8.2 performed feature tracking from the reaching videos.

DeepLabCut labeling utilized a marker set that was chosen to track the position of fingers, hands, cricket, and features of the reaching platform. We trained the pre-trained ResNet50 network supplied by DeepLabCut using 574 manually labeled, manually selected frames of video. The set of frames was selected to fully characterize the diversity of marmosets, camera angles, hand positions, and targets of our reaching video data. The final model was trained for 800k iterations and used the imgaug augmentation pack. Deeplabcut was deployed through Anipose, which generated unfiltered 2D tracking files using DeepLabCut algorithms. In some later recordings, an unexpected minor shift in the top-view camera position **(Figure 1F)** offset the accuracy of the original triangulation calibration. To correct for this, we re-aligned the 2D DeepLabCut tracking data from each reach video segment using a linear transformation matrix generated by comparing the arena feature markers **(Figure 1D)** taken from the original calibration videos to the arena feature markers of each reaching video. We measured the distance each point was moved to confirm that transformation did not introduce unintended scaling artifacts. These steps were performed using custom Matlab scripts after converting tracking data files to .csv files. All tracked points were transformed accordingly, re-packaged into .hd5 files using custom Python scripts, and then triangulated into 3D space using Anipose.

### Reaching Kinematic Data Preparation

Manual reach scoring provided metadata for each reach, which guided the identification of video segments with hand extension and target motion position data. Position estimates with less than a .7 estimation likelihood were excluded from analyses, which is a metric provided by DeepLabCut indicating label accuracy probability. Hand and cricket tracking points were filtered using a 5 sample Gaussian smoothing window. We applied a 180-degree rotation of right-handed reaches and the cricket position about the reach platform midline to reduce any potential effects of any handedness-related directionality biases.

We defined a reach as the first tracked appearance of the central hand point beyond the reach aperture plane to the last tracked point of the cricket. Cricket tracking at the very end of reaches, during deacceleration and grasp, was typically lost due to occlusion by the fingers or hand. Windows of reaching data were defined by viewing videos with overlayed tracked points and marking the beginning and end frame number. These frame number values were used to refer directly to the .csv tracking data files in custom Matlab analysis scripts. Reaches for crickets that moved greater than a 0.5 cm distance during the reach and moved continuously were included in analyses. Crickets that moved this distance by a jump or that ceased motion during the reach were not included in the data set, nor were crickets that moved less than 0.5cm in total.

### Grasp Cluster Analyses

Reaches were selected by visualizing all the tracked points of each of the 78 selected reaches with moving crickets and excluding 14 reaches with poor knuckle and finger tracking. Each reach was interpolated to a 51-sample vector where each point is equal to the mean distance between the pointer and middle knuckle and middle and ring knuckle markers. Knuckle markers were chosen because they were better tracked and less subject to perturbation by the cricket retaining wall. Each finger separation vector was z scored and the population was subjected to a kmeans cluster analysis. The maximum number of clusters was set manually based on testing 2-6 clusters. We repeated clustering using different initial centroids 10 times. The results of this process were nearly identical to a hierarchical wavelet-based clustering process performed using the mdwtcluster function provided by Matlab.

### Pursuit Models and Velocity Decomposition

To investigate the degree to which marmoset reaching adheres to steering guidance laws such as pure pursuit and proportional navigation, we adopted Matlab analysis scripts used to investigate hawk pursuit of moving targets(Brighton & Taylor, 2019). These scripts provided the framework to simulate marmoset reaches by applying the sample-by-sample steering commands while matching the magnitude of each step to the measured actual hand speed. Simulation fits to tracked hand data were measured using the R-squared between the simulated and actual hand velocity across each reach, then pooled across all reaches. To set initial conditions for simulated reaches, we matched simulated hand velocity in the first 75ms of the reach to the mean of the actual hand velocity averaged over the same period. Following Brighton and Taylor(Brighton & Taylor, 2019), the proportional navigation guidance law utilized by these scripts is defined as

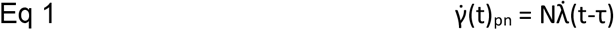

Where 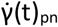 is turning rate, N is an experimentally derived gain constant, 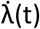 is the line of sight rate, t is time, and τ is a delay parameter. Proportional pursuit is defined as

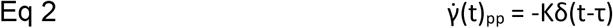

Where 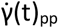 is turning rate, K is an experimentally derived gain constant, δ is the range vector angle with respect to the hand’s velocity, t is time, and τ is a delay parameter. A mixed strategy is defined as the sum of the two above terms as follows:

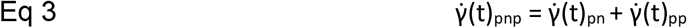

Model parameters for gain were set by maximizing the total R-squared for hand velocity against model hand velocity. All reach trials (N=78) were included to fit a gain parameter that was fixed across trials. Confidence intervals on the R-squared performance were computed from the standard deviation of trial-by-trial R-squared performance.

Proportional navigation strategies yield parallel pursuit, which can be described as matching velocity orthogonal to the line of sight, or range vector. To investigate the behavior of range-vector-relative decomposed velocities in the above steering model frameworks, target and hand velocity vectors were decomposed into orthogonal velocity components relative to a range vector, which connects hand position to cricket position. We refer to the velocity component parallel to the range vector as *direct* velocity. We refer to the velocity component orthogonal to the range vector as *lateral* velocity. This distinction is important due to the typical movement of crickets along the perimeter of the reaching platform. When cricket velocity is decomposed in this way, the majority of cricket velocity is contained within the *lateral* velocity component. A summary of this decomposition is provided in **Figure 3D**.

A proportional navigation analysis was conducted according to methods used in dragonfly pursuit(Mischiati et al., 2015). Range vectors were calculated as a subtraction of hand position from cricket position. The time derivative of the series of range vectors was calculated and correlated with the range vector itself using cosine vector correlation. Proportional navigation predicts a negative correlation of −1 between the range vector and its derivative.

### Predictive Modeling and Reaching Analyses

We examined if sudden increases in cricket velocity during reaching led to corresponding increases in hand velocity. To identify increases in cricket velocity we filtered the lateral speed of the cricket (absolute value of lateral velocity) using a difference filter that was defined by a Gaussian kernel (σ = 20ms) with negative amplitude at negative lags and positive amplitude as positive lags. This filter produces the largest values for epochs where low speed is directly followed by an increase in speed, regardless of whether speed increase is due to the cricket moving to the left or right. We used a threshold to flag the top 5% of the filtered values and identify peaks, ensuring that no two peaks occurred within 100 ms of each other. From 78 total reach trials we flagged 68 events. We averaged the lateral cricket velocity and hand velocity traces from −100 to 150 ms time-locked around these events (**Figure 4A**). Results were similar if we relaxed our acceleration threshold to include the top 10% or 20% of events (not shown).

Decomposed hand and cricket velocity components were used to investigate the predictive influence of cricket velocity on hand velocity. We first tested a linear prediction in which the lateral hand velocity was predicted by a weighted linear sum of the lateral cricket velocity history. The linear predictor equations was defined as

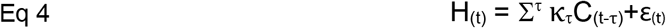

where H_(t)_ represents lateral hand velocity at time t, C is the lateral cricket velocity at time t minus time delay τ and κ_τ_ is a linear prediction coefficient at this time delay. To reduce the number of parameters we down-sampled the velocity traces from 240 to 80 hertz, giving 12.5ms steps. We tested a range of τ values from −25 to 150ms. We defined the range vector used to separate direct and lateral velocities as the vector from the hand to cricket at the start of the reach (when the hand first entered the arena). We considered the case of updating the range vector at each delay during the reach as in the full auto-regressive model discussed below and settled upon this approach for the sake of simplicity.

A recent study examined a pursuit strategy embedded in a 2D coordinate system with a term incorporating prey velocity to test for the role of prediction (Yoo et al., 2020). To apply the same model, we employed an auto-regressive linear regression to fit parameters in Eq 5 shown below which includes a momentum term for hand velocity, and a term based on the predicted range vector. The parameters for visuo-motor delay, τ_d,_ and a prediction time constant, τ_p,_ were fit to optimize the match to actual hand velocity. This analysis was evaluated using the Matlab *mvregress* function. Concatenated velocity and range vector magnitude values from each reach were supplied to the regression function. Time is defined by frames of video, each of which captures 1/240 second, or 4.2ms of behavior.

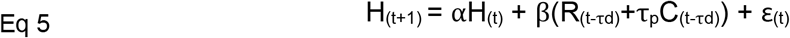

In the above Equation 5, H_(t+1)_ represents hand velocity one frame of video into the future (t+1) relative to the current H_(t)_ velocity at time t. The velocity of the cricket C at a delayed time, t-τ_d_, is multiplied by the prediction time constant, τ_p,_ to project cricket position into the future or past. We also include a variable representing the range vector from hand to the cricket position as R. This vector connects hand position at time t and cricket position at t-τ_d_. The sequence of operations to generate these variables was to first temporally offset tracked cricket position values according to t-τ_d_. Velocities and range vectors were calculated. Finally cricket velocities were multiplied by τ_p_. The predictor variables are weighted by a set of coefficients and, respectively. All terms in this equation are composed of cartesian x and y constituents and thus are bivariate. These data were submitted to a least-square regression where ε _(t)_ represents the residual error.

We varied the values of τ_p_ and τ_d_ to determine the effects of visuomotor delay and prediction on model fit. The parameter τ_d_ was varied from 0 to 35 frames of video (equivalent to 0ms to 146ms) such that time shifted cricket position values were aligned to present hand values. Velocities and range vectors were calculated after this offset was applied. Following this we multiplied cricket velocity by a range of τ_p_ values ranging from −60 to 60 frames of video (−250ms to 250ms). To quantitatively assess the influence of cricket velocity on model fit and to subsequently find offsets with significant cricket velocity influence, we compared the Akaike Information Criterion (AIC) (Akaike, 1973) of the model described in Eq5 to analogous models that exclude the range vector and cricket velocity predictor terms R and C. This was performed for each τ_p_ and τ_d_ in the above range. When the AIC value of one model is more negative than another, it can be interpreted to have a better fit. More negative AIC values in the cricket inclusion models would suggest that prediction based on cricket velocity is a significant contributing factor to reach kinematics. We plotted the AIC difference values and used a jackknife approach (leaving ∼8 of 78 reaches out for 10 jackknife iterations) to estimate jackknife AIC mean and calculate standard error.

To compare the predictive pursuit model against steering models we applied the simulation approach discussed above in Eq. 1-3 and compared its R squared performance for matching hand velocity against those models. All features of the simulated reaches were applied as before for the steering models with the guidance law described by Eq. 6 as

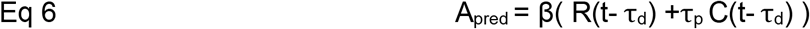

where A_pred_ is acceleration applied to hand velocity, β is an experimentally derived gain constant, R(t-τ_d_) is the range vector and C(t-τ_d_) is the cricket velocity, both at delay τ_d_, and τ_p_ is the prediction time constant. All 78 trials were included to maximize the total R-squared fit across trials in fitting β and τ_p_ parameters for each tested visuo-motor delay τ_d_. Confidence intervals on the R-squared performance were computed from the standard deviation of trial-by-trial R-squared values with statistical tests comparing trial R-squared between models (Sign Rank tests).

## Supplemental Discussion

The benefits of utilizing naturalistic behavior for neuroscience research are tempered by experimental variability. However, modeling approaches provide a tractable alternative to traditional trial averaging for addressing this variability issue. Naturalistic behavior can tap into more dynamic aspects of visually-guided reaching including prediction. Furthermore, this approach presents animals with tasks that they are already accustomed to performing. As a clear example, our research was conducted with none of the extensive animal training effort that is required for traditional vision task-related experiments with non-human primates. However, both the subject and the experimental environment (e.g., crickets) have larger intrinsic variability, which increases the difficulty of collecting trials that meet inclusion criteria and performing traditional averaging over repeated trials. We circumvented the lack of identical repeats by performing model fitting, which made it possible to determine the influence of visuomotor delay prediction on behavior. Future research could benefit from approaches that leverage the variability of movements to examine the sensitivity of individual reach trajectories to external or internal disturbances such as unexpected cricket motion or gaze shifts. Finally, while the reaching task utilized here is less constrained than traditional reaching tasks, it still imposes some physical constraints on reaching. This may have precluded more diverse strategies including direct capture using the mouth, which has been found in the wild(Ngo et al., 2022; Schiel, Souto, Huber, & Bezerra, 2010). Nonetheless, the experiments and analyses described here represent an intermediate step towards growing efforts to achieve more naturalistic behavior in laboratory settings (Datta, Anderson, Branson, Perona, & Leifer, 2019).

Previous model-based approaches have shown that primates utilize simple physical parameters to predictively pursue an evading virtual target with a joystick (Yoo et al., 2020). A linear prediction term in the model employed by Yoo and colleagues allowed them to estimate a visuo-motor delay in their task. Similar to our result, they find evidence for a predictive strategy, although their estimate of visuo-motor delay differs substantially from the value reported here. This can likely be attributed to differences in their task design because movements of the cursor on the screen were limited by the speed allowed under joystick control and momentum terms therein, with overall much slower timescales for target and cursor speeds on the order of seconds. By contrast the natural dynamics of marmoset reaches unfold typically in under 200ms. The approach taken in our study emphasizes the dynamics of natural behavior and motor control, which enables interrogation of features of the visuomotor system within the natural timescales of visually-guided reaching and target pursuit.

The derived estimate of marmoset visuomotor delay is consistent with experimentally derived estimates in humans given the quicker processing and neural transduction times of smaller primates. A widely studied paradigm that interrogates the on-line visuomotor response delay is the double step task. In this task, subjects reach for an initial target, which is extinguished and replaced by a second target during the flight of the hand. The limited number of studies with macaques that have used double-step reaching tasks to probe online correction have found substantially longer motor delays than our estimation, around 200-300ms(Philippe S. Archambault et al., 2009; Battaglia-Mayer et al., 2013; Dickey et al., 2013; Georgopoulos et al., 1981). Of note, the redirected reaches are typically preceded by a saccadic eye movement at 100-200ms latency to the new jumped location followed by changes in the hand trajectory at 50-100ms (Archambault et al, 2015). Further, the jumps of targets used in non-human primate studies have often been much larger than those in human studies, which could impose additional delays in redirecting attention before movement adjustment. On-line corrective reaction times, which are analogous to an on-line visuomotor response delay, have been reported in humans from 125ms to > 200ms (Brenner & Smeets, 1997; Brouwer, Brenner, & Smeets, 2002; Day & Lyon, 2000; De Brouwer & Spering, 2021; Prablanc & Martin, 1992; Reichenbach, Thielscher, Peer, Bulthoff, & Bresciani, 2009). This range is likely due to differences in task parameters (eg. pointing with a finger, holding a rod while reaching, using a pantograph, etc.), measurement methods (eg. touch plates, receiver operating characteristic analyses, scored deviation from control, etc.), and the influence of separate motor control pathways with different latencies (eg. tectospinal versus corticospinal). If we take the bottom line from one of these human experiments that derives delay from a deviation in hand kinematics (Day & Lyon, 2000), which is 125ms, and compare that to our estimate of marmoset reaching delay, which is 80ms, we are within range of a human/marmoset difference seen in a separate, but related comparison of human and marmoset kinematics. The ocular following response measures the automatic movement of the eye when presented with coherently moving dot patches. In a study of both humans and marmosets, humans demonstrate a 125ms delay in ocular following response while marmosets demonstrate a 75ms delay, which aligns well with our findings (Cloherty et al., 2020). While the motion of the crickets in this reaching paradigm is not controlled in the same way as in double step psychophysics experiments, it does demand adjustments in reaching to account for its motion and thus provides sufficient variation to estimate visuomotor delay through model fitting.

While insect hunting is effective for tests of marmoset reaching and has been quantified in natural behavior recently in wild marmosets(Ngo et al., 2022), we do find some limitations using this approach that could be easily addressed in future laboratory studies. In particular, for our study the number of reaching movements in which the cricket moved during the reach were relatively small in proportion (∼30% of reaches). Further, the number of reaching movements that we could obtain in any single behavioral session was limited to roughly 15 crickets in a day, more than enough to satiate a marmoset. Given that only a small fraction of those movements involves cricket motion during the reach, we obtained relatively few dynamic reaches per day in any given animal. This limitation of data quantity motivated our pooled approach across 10 marmosets to analyses. By pooling movements across animals and days we were able to estimate the visuomotor delay, but it would be difficult to employ this same paradigm to test causal manipulations of reaching behavior measurable within a single behavioral session. Future work may seek an intermediate solution without live crickets, such as reaching to small food treats that are moved on a robotic arm, in which the movement of the target can be controlled under online feedback during the reach.

We found that DeepLabCut-powered tracking provided reliable descriptions of the trajectory of the hand supporting the reliability reported in a previous study(Moore, Walker, MacLean, & Hatsopoulos, 2022). DeepLabCut affords a key advantage for studying natural movements over other methods that affix optical tracking markers to hands and arms, which may impose constraints on animal reaching and require additional time and care to install. We found that GoPro cameras were well suited for data collection in these reaching tasks due to their ability to capture high frame rate video with user-set shutter speeds that reduce motion blur. While the arrangement of cameras was optimized for a 3-D rendering of the top-down view of the hand, occlusion of the fingers was common due to the wide variability of reach trajectories, the pronation of the hand, and the tendency of the fingers to make contact with the cricket retaining wall. Regardless, it is impressive that DLC could provide tracking of features with such small degrees of error and could discriminate between the right and left hands without confusing their features. One limitation of our current tracking approach is that it is difficult to track body parts that are occluded by featureless fur such as the shoulder or elbow. Future studies could characterize the detailed interaction between shoulder and elbow movement in positioning the final hand position during motor control, as well as the role of gaze in guiding reaching movements. Finally, we found it difficult to achieve low-noise 3D depictions of reaching and grasping and thus used 2D tracking for our analyses. We feel this problem is likely solved by additional camera angles to both generate more camera coverage overlap and to better cover areas that were poorly captured by the set of 3 cameras used in this study.

## Supplemental Figures

**Supplementary Figure 1.**
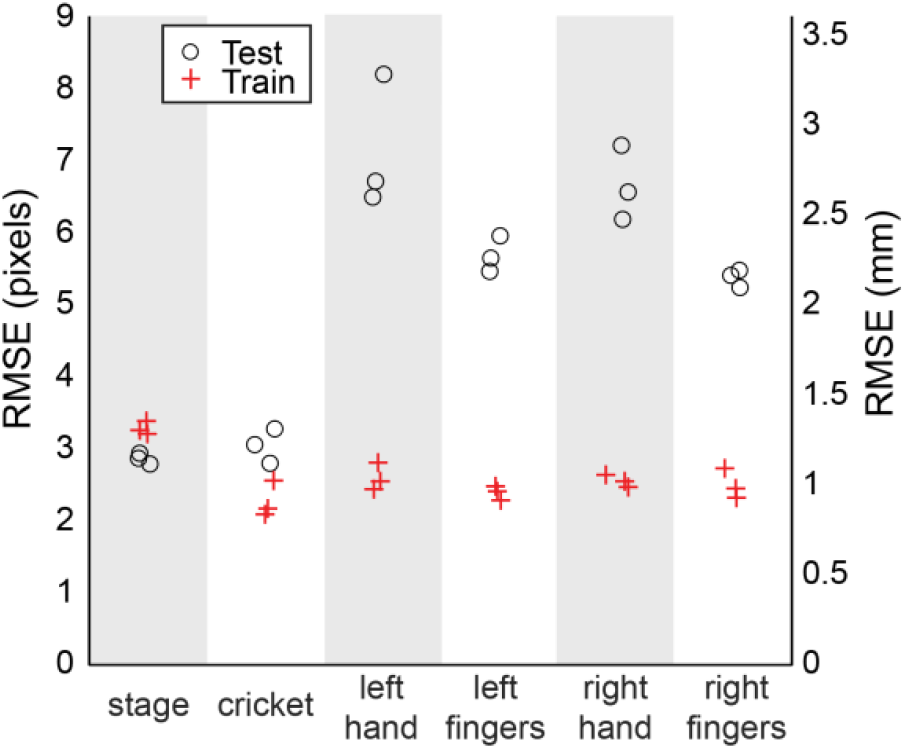
Tracking accuracy in DeepLabCut video analysis: Three DeepLabCut networks were trained to show tracking errors from the testing and training sets in RMSE quantified in pixels and converted to mm using the pixel to mm conversion respective to the above-view camera (Figure 1F). The label ‘stage’ on the x-axis refers to the reach platform stage and corresponding arena points (Figure 1D). Fingers refers to the average of both the fingertip and knuckle points, which equals 10 points for each hand. The average RMSE values depict the stability of the training approach over 3 different training iterations. Each point depicts the labeling error of a model trained with a different “shuffle” of the test and training data. The model used for all tracking analyses performed with a testing RMSE of 2.79 pixels for the stage and arena markers, 3.28px for the cricket marker, 8.21px for the left hand, 5.1px for the left fingertips, 5.9px for the left knuckles, 6.2px for the right hand, 5.8px for the right fingertips, and 5.0px for the right knuckles. For comparison, the width of the wrist is typically around ∼1.5cm, which equates to 37px. The length of the cricket ranged from ∼1.25cm to ∼3cm. We utilize the hand marker for reach modeling and correlation analyses because of its tracking reliability across all three camera views and its reduced susceptibility to occlusion-related tracking drops, which were more common in the finger markers.

**Supplementary Figure 2.**
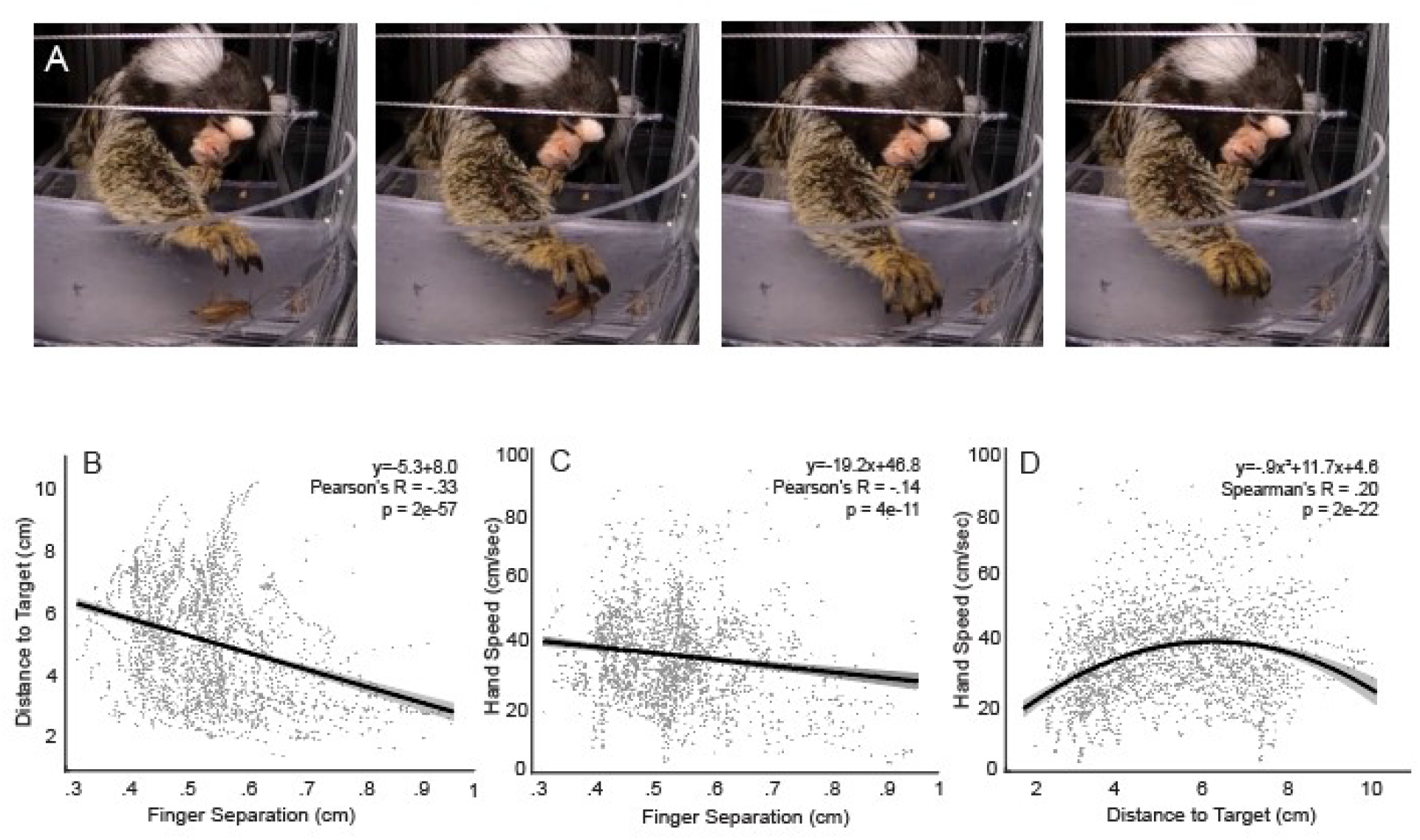
Further grasp characterization in relation to hand speed and distance. **A:** Grasp visualization from a low camera angle to show parallel hand orientation and simultaneous finger closure. **B-D**: Scatter plots illustrate the relationships between distance to target, hand speed, and finger separation across reaching movements. Points show a 1/4 subsample of all tracked points from the population of reaches. Each plot is fit with a line (B,C) or 2^nd^ order polynomial (D), which is shown with a 95% confidence interval. **B:** Across many reaches the finger separation increases with proximity to target. **C:** Finger separation increases with decreased hand speed. **D:** Hand speeds are lower when at greater distances from target and when in close proximity to the target.

